# Linking bacterial tetrabromopyrrole biosynthesis to coral metamorphosis

**DOI:** 10.1101/2023.05.08.539906

**Authors:** Amanda T. Alker, Morgan V. Farrell, Alyssa M. Demko, Trevor N. Purdy, Sanjoy Adak, Bradley S. Moore, Jennifer M. Sneed, Valerie J. Paul, Nicholas J. Shikuma

**Affiliations:** Department of Biology and Viral Information Institute, San Diego State University, San Diego, California 92182 USA; Smithsonian Marine Station, Ft. Pierce, FL 34949, USA; Center for Marine Biotechnology and Biomedicine, Scripps Institution of Oceanography, University of California, San Diego, La Jolla, CA, 92093

## Abstract

An important factor dictating coral fitness is the quality of bacteria associated with corals and coral reefs. One way that bacteria benefit corals is by stimulating the larval to juvenile life cycle transition of settlement and metamorphosis. Tetrabromopyrrole (TBP) is a small molecule produced by bacteria that stimulates metamorphosis in a range of coral species. A standing debate remains, however, about whether TBP biosynthesis from live *Pseudoalteromonas* bacteria is the primary stimulant of coral metamorphosis. In this study, we create a *Pseudoalteromonas* sp. PS5 mutant lacking the TBP brominase gene, *bmp2*. Using this mutant, we confirm that the *bmp2* gene is critical for TBP biosynthesis in *Pseudoalteromonas* sp. PS5. Mutation of this gene ablates the bacterium’s ability in live cultures to stimulate the metamorphosis of the stony coral *Porites astreoides*. We further demonstrate that expression of TBP biosynthesis genes is strongest in stationary and biofilm modes of growth, where *Pseudoalteromonas* sp. PS5 might exist within surface-attached biofilms on the sea floor. Finally, we create a modular transposon plasmid for genomic integration and fluorescent labeling of *Pseudoalteromonas* sp. PS5 cells. Our results functionally link a TBP biosynthesis gene from live bacteria to a morphogenic effect in corals. The genetic techniques established here provide new tools to explore coral-bacteria interactions and could help to inform future decisions about utilizing marine bacteria or their products for restoring degraded coral reefs.

## MAIN TEXT

Some marine bacteria stimulate the early life-cycle transition from larval to juvenile phases in corals (i.e., metamorphosis). These bacteria could be used to promote coral larval recruitment in the wild, cultivate corals for reseeding degraded reefs or the aquarium trade, or help test basic science questions in the laboratory [1–3]. Single-species biofilms of *Pseudoalteromonas* bacteria, such as *Pseudoalteromonas* sp. strain PS5, have been shown to stimulate coral metamorphosis [2, 4, 5]. Moreover, the compound tetrabromopyrrole (TBP), extracted from *Pseudoalteromonas* bacteria or chemically synthesized, robustly promotes the metamorphosis of diverse coral species with and without attachment [2, 5–7] (Figure 1A). However, an open question remains about whether live *Pseudoalteromonas* bacteria stimulate coral larval metamorphosis solely by producing TBP, or whether these *Pseudoalteromonas* strains produce yet unknown products that confer part or all of the stimulatory activity [6, 8].

**Figure 1.**
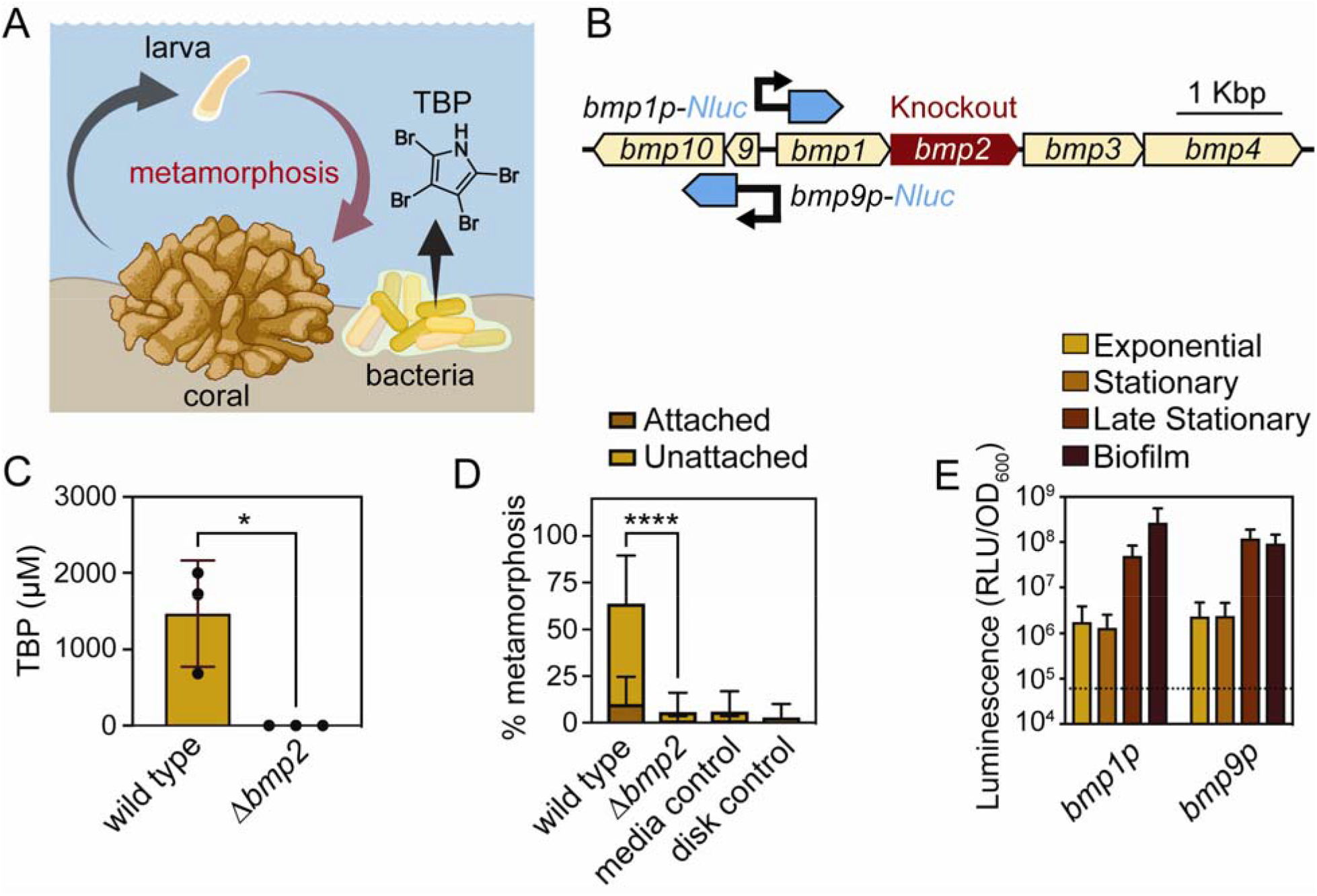
TBP biosynthesis in *Pseudoalteromonas* sp. PS5 induces metamorphosis of the stony coral *Porites astreoides*. (A) A model of TBP biosynthesis in *Pseudoalteromonas* sp. PS5 and its ability to induce coral metamorphosis. (B) Synteny of the *bmp* gene cluster in *Pseudoalteromonas* sp. PS5 (Genbank accession KR011923) created with EasyFig (V1.4.4) [9]. The 5’UTR for both *bmp1* and *bmp9* were cloned and fused to a *Nanoluc* (*Nluc*) reporter shown in blue. The *bmp2* brominase is highlighted in red (C) LC-MS/MS of *Pseudoalteromonas* sp. PS5 wild type and *bmp2* knockout quantifying tetrabromopyrrole production (one-tailed Mann-Whitney test, P = 0.05). Error bars show standard deviations. N = 3 extractions from replicate 5 mL liquid cultures. (D) Metamorphosis biofilm assays (%) with *Porites astreoides* larvae (10 larvae/ dish) in response to *Pseudoalteromonas* sp. PS5 wild type and Δ*bmp2* strains. Control treatments include disks incubated in sterile Marine Broth media or filtered seawater (FSW). N=25 total dishes, including 3 experiments spanning two collection years. Combined morphogenesis significance is shown (Dunn’s multiple comparison, Adjusted P < 0.0001). (E) Luciferase assays of *bmp1* and *bmp9* promoter expression under different modes of growth reported in relative luminescence units normalized by the optical density (RLU/ OD_600_) and plotted on the Log_10_ scale. The 5’UTR of *bmp1* and *bmp9* were compared against the negative control (*Pseudoalteromonas* sp. PS5 cells expressing a non-luminescent plasmid) as represented by the dotted line (Y = 60,422 RLU/ OD_600_). Error bars show standard deviation of the mean. N = 4 biological replicates.

In *Pseudoalteromonas* sp. PS5 and phylogenetically related strains, TBP biosynthesis from L-proline is performed by three enzymes and a carrier protein encoded by the brominated marine pyrroles/phenols (*bmp*) gene cluster [10], specifically genes *bmp1-4* [11] (Figure 1B). We thus sought to create a TBP deletion mutant in *Pseudoalteromonas* sp. PS5 to query coral larval metamorphosis by genetically inactivating the *bmp2* gene, which codes for the brominase that installs all four bromine atoms in TBP [11]. We determined that *Pseudoalteromonas* sp. PS5 is amenable to genetic manipulation by conjugation using a broad-host-range *gfp* reporter plasmid from the Marine Modification Kit (MMK) plasmid system [12]. We generated a *Pseudoalteromonas* sp. PS5 mutant with an in-frame deletion of the *bmp2* gene (Δ*bmp2*) following previously established methods for double-homologous recombination [13] (Figure 1B). This mutation, as anticipated, resulted in the complete loss of TBP production in the Δ*bmp2* strain in stark contrast to the wild-type culture that readily produces TBP (1468 ± 400.8 μM TBP, P = 0.05, one-tailed Mann-Whitney test) (Figure 1C). These results show that the *bmp2* gene is required for TBP production in *Pseudoalteromonas* sp. PS5 under the conditions tested.

We next tested whether bacteria lacking the *bmp2* gene were able to stimulate the metamorphosis of the Caribbean coral, *Porites astreoides*, which has been shown previously to undergo metamorphosis in response to *Pseudoalteromonas* sp. PS5 and TBP [2]. When exposed to biofilms of wild type *Pseudoalteromonas* sp. PS5, we observed the metamorphosis of coral larvae consistent with previous findings [2, 4–6], both attached to the substrate (11.6% ± 7.2, Adjusted P = 0.0002, Dunn’s multiple comparisons test) and unattached (55% ± 13.1, Adjusted P < 0.0001, Dunn’s multiple comparisons test) (Figure 1D). In contrast, the treatment with biofilms of the Δ*bmp2* strain resulted in a reduced ability to stimulate the metamorphosis of coral larvae compared to the wild type strain (Adjusted P < 0.0001, Dunn’s multiple comparisons test) (Figure 1D). We observed some attached (0.6% ± 2.9) and unattached (5.1% ± 10.4) metamorphosis in the Δ*bmp2* treatment at comparable to the background rates in the media and disk controls. Our results demonstrate that the effect of *Pseudoalteromonas* sp. PS5 on coral metamorphosis is primarily due to the production of TBP.

We then questioned whether different growth conditions affect the expression of the *bmp* genes in *Pseudoalteromonas* sp. PS5. To this end, we cloned the *bmp1* and *bmp9* promoters, fused them with a *NanoLuciferase* (*NLuc*) reporter gene, and conjugated the resulting plasmids into *Pseudoalteromonas* sp. PS5 (Figure 1B) [12]. When measured in exponential, early or late stationary (Figure S1), or biofilm growth phases, the *bmp1* promoter displayed a 203-fold activity range, with the highest expression in late stationary and biofilm phases (Figure 1E). The *bmp9* promoter followed similar expression profiles, suggesting that the gene cluster may be co-regulated (Figure 1E). We also tested broad-host-range promoters, which displayed at least a 457-fold increase in expression compared to assay background across all tested conditions (Figure S2). Our results suggest that the expression of TBP biosynthesis genes is strongest when bacteria exist in a slow-growth state when *Pseudoalteromonas* sp. PS5 might occur within surface-attached biofilms on the sea floor.

*Pseudoalteromonas* species are known to associate with marine eukaryotes and produce interesting antimicrobial metabolites [14], yet the study of these host-microbe and microbe-microbe interactions remains challenging because of a lack of available tools for their genetic manipulation. We therefore developed an integrative Tn10 transposon for use in *Pseudoalteromonas* sp. PS5, which is compatible with existing modular genetic toolkit parts [12, 15, 16] (Figure S3). With the Tn10 transposon, we generated *Pseudoalteromonas* sp. PS5 with *gfp*-tags integrated into the genome, which were confirmed by whole genome sequencing to identify genomic insertion loci (Table S1).

In this work, the genetic engineering of a marine microbe enabled us to test a standing question about the role of TBP in coral metamorphosis. Our results represent the first functional characterization of a gene in a marine bacterium conveying a beneficial effect in corals. Previous studies suggest that *Pseudoalteromonas* species may not be present in ecologically relevant concentrations that would stimulate coral metamorphosis [6]. However, TBP could be considered as a metamorphosis stimulant tool, for example, to enhance coral settlement *ex situ* for outplanting on coral reefs. The strains and genetic tools developed in this study could be helpful for dissecting how TBP stimulates metamorphosis with and without attachment in corals [17] and for studying TBP’s effects on eukaryotic cell physiology [18–21]. Beyond TBP, our results demonstrate how genetic methods can help characterize genes and gene products from bacteria in the context of marine microbial interactions, providing new techniques to interrogate the microbial ecology of *Pseudoalteromonas* spp. We hope this work will serve as a model for elucidating function in bacteria-coral interactions and will inform the use of bacteria for coral reef restoration [22].

## Acknowledgements

Corals were collected under FKNMS-2019-24 permit from the Florida Keys National Marine Sanctuary. We would like to thank SECORE International and Florida Department of Environmental Protection for support and personnel provided for coral larval collection. We would like to thank Erich Bartels, Cory Walter, Joe Kuehl, and Samantha Simpson, for collecting and returning the *Porites astreoides* colonies and Zach Ferris, Natalie Danek and Samantha Scheibler for helping with larval collection over the 2021-2022 larval collection seasons at the Mote Marine Laboratory’s Elizabeth Moore International Center for Coral Reef Research and Restoration in Summerland Key, Florida. We thank Tommy Demarco, Carle Dugan, Yesmarie de la Flor and Kelly Pitts for help setting up and scoring the coral metamorphosis assays. We would also like to thank Dr. Blake Ushijima, Dr. Kristen Marhaver and Dr. Raphael Ritson-Williams for guidance and helpful discussions about coral metamorphosis and microbiology. Finally, thank you to the Shikuma Lab members, including Dr. Tiffany Dunbar, Dr. Kyle Malter, Dr. Kate Nesbit, Emily Darin and Andy Fedoriouk for their feedback and support with cloning, imaging, and editing the manuscript. Schematic figures were created in part using Biorender.com.

This work was supported by the National Science Foundation (2017232404, A.T.A.; 1942251, N.J.S.; OCE-1837116, B.S.M.), the Gordon and Betty Moore Foundation (GBMF9344 to N.J.S.; https://doi.org/10.37807/GBMF9344), Office of Naval Research (N00014-20-1-2120 to N.J.S.), the National Institutes of Health, (R35GM146722 to N.J.S.; R01ES030316 to B.S.M.) and the Alfred P. Sloan Foundation, Sloan Research Fellowship (N.J.S.). A.T.A. and N.J.S. are coinventors on provisional U.S. patent application Serial number 63/323,653, entitled “Genetic Engineering of Marine Bacteria for Biomaterial Production, Probiotic Use in Aquaculture and Marine Environmental Restoration” and assigned to San Diego State University Research Foundation.

