## Supplemental Methods, Figures, Tables for "Linking bacterial tetrabromopyrrole biosynthesis to coral metamorphosis"

**SUPPLEMENTARY MATERIALS AND METHODS**

**Bacterial strains and growth conditions**

The strains and plasmids used in this study are listed in Tables S2 and S3. The strain *Pseudoalteromonas* sp. PS5 was first isolated from the surface of *Paragoniolithon solubile*, a species of crustose coralline algae collected from a fringing reef near Looe Key, FL [1]. *Pseudoalteromonas* sp. PS5 was cultured with natural seawater tryptone media NSWT (1 L 0.2 µm filtered natural seawater, 2.5 g tryptone, 1.5 g yeast, 1.5 mL glycerol) or Marine Broth (2216 Difco) media and incubated between 25-28 °C. *Escherichia coli* cultures were grown in LB media at 37 °C. All liquid cultures were inoculated with a single colony and incubated between 14-18 hours while shaking at 200 rpm unless otherwise indicated. Plasmids were selected and maintained on LB kanamycin 100 µg $mL^{-1}$ or LB kanamycin/ampicillin 100 µg $mL^{-1}$.

**Generation of *bmp2* knockout strain**

Primers used to generate strains in this study are listed in Table S4.

The *bmp* gene cluster (Genbank accession KR011923) was identified by Blastn from the *Pseudoalteromonas* sp. PS5 genome [2]. The in-frame deletion of *bmp2* was generated following a previously published protocol [3]. Briefly, Gibson primers were ordered from integrated DNA technologies (IDT) and were designed to amplify 1400 base pair (bp) homology arms up and downstream of the *bmp2* gene in *Pseudoalteromonas* sp. PS5. The homology arms were amplified using a high-fidelity DNA polymerase (Primestar, Takara) and the resulting fragments were purified using a DNA Clean and Concentrator Kit (Zymo Research). The suicide vector pCVD443 [4] was digested with SphI, XbaI, and SacI. To assemble the digested plasmid and the

PCR products, a three fragment Gibson Assembly was performed using the NEBuilder HiFi DNA Assembly Master Mix at a ratio of 2:1 for inserts: backbone vector. Resulting assemblies were diluted and electroporated into SM10*pir* electrocompetent cells and selections were performed on LB ampicillin 100 µg $mL^{-1}$. Positive clones containing a band around 3000 bp were cultured and miniprepped using the Zyppy DNA Miniprep Kit (Zymo Research). Minipreps of positive clones were confirmed by Sanger sequencing (Eton Biosciences). The pCVD443_PS5Δ*bmp2* plasmid was conjugated with *Pseudoalteromonas* sp. PS5 according to a previously published double homologous recombination protocol [3]. Selections in *Pseudoalteromonas* sp. PS5 were performed on NSWT streptomycin/kanamycin 200 µg $mL^{-1}$ and counter selections were performed on NSWT + 10% sucrose.

**Cloning and modular Golden Gate Assembly**

Native promoters (pMMK204/205) and the *bmp2* gene (pMMK301) were PCR amplified from *Pseudoalteromonas* sp. PS5 and Gibson assembled into linearized Type 2 and Type 3 plasmid parts containing the appropriate 4 bp overhangs. Modular part plasmids compatible with the BTK, YTK, MMK system were miniprepped (Zyppy Plasmid Miniprep) to generate Stage 1 assembled Golden Gate plasmids via the BsaI-HF V2 Type IIS restriction enzyme (NEB) and ligation with T4 Ligase (Promega) following previously described methods [5, 6]. Thermocycler conditions for the assembly were used as follows: 37 ºC for 5 minutes, 16 ºC for 5 minutes, cycles 1-2 repeated 30 times, followed by 37 ºC for 10 minutes and 80 ºC for 10 minutes. 2 µL of assembly was electroporated directly into S17-1*pir* electrocompetent cells and then clones were PCR confirmed. All conjugations were performed using MFD*pir* auxotrophic donor cells and followed a previously established protocol [6].

**Growth curve of *Pseudoalteromonas* sp. PS5 strains**

*Pseudoalteromonas* sp. PS5 wild type and knockout *bmp2* strains were grown on Marine Broth (MB) agar plates and incubated overnight at 25 °C. PS5 strains expressing plasmids were grown on MB agar plates with 300 µg $mL^{-1}$ of kanamycin and grown overnight at 25 °C. Single colonies were picked and inoculated into 5 mL of MB liquid media with respective antibiotics listed above. Two biological replicate cultures were inoculated for each strain by picking different colonies from the agar plate and inoculating separate 5 mL cultures. Cultures were incubated at 25 °C for 16 hours shaking at 200 rpm. From the initial cultures a subculture was created by performing a 1:100 dilution into the subculture. The subculture consisted of 25 mL of MB liquid media and 250 µL of original culture along with the respective antibiotics in 125 mL flasks. Subcultures were incubated at 25 °C shaking at 200 rpm throughout the growth curve experiment. Optical density (OD) at a wavelength of 600 nm was measured from the subculture every hour for the first 10 hours and then a final measurement was taken at 24 hours. Plotted is the average $OD_{600}$ from 2 biological replicates reported on a $log_{10}$ scale. A Gompertz non-linear regression model was applied to the wild type (K = 0.96 hours, Ym = 2.583 $OD_{600}$) and Δ*bmp2* strains (K = 0.58 hours, Ym = 3.095 $OD_{600}$).

**Luciferase assay**

The luciferase assay was performed using a previously established protocol [6] with the following adjustments. *Pseudoalteromonas* sp. PS5 containing plasmids with either broad-host-range or native promoters driving a NanoLuciferase reporter were grown on Marine Broth with 300 µg $mL^{-1}$ of kanamycin, in either liquid culture or agar plate (referred to the biofilm treatment for this assay). Liquid cultures were monitored by measuring $OD_{600}$ to delineate growth phases with exponential being defined as an $OD_{600}$ of 0.35-0.67, early stationary $OD_{600}$ 1.0-1.45, and late stationary $OD_{600}$ 2.38-2.54 (Figure S1). Nanoluciferase reactions were performed through serial dilution in 10-fold steps until a consistent reaction rate was found. For exponential cultures, *Pseudoalteromonas* sp. PS5 broad-host-range promoters (PA3, Ptac, CP25) were diluted by 1:10 or 1:1,000, whereas native promoters (*bmp1* and *bmp9*) were diluted by 1:100. For stationary phase cultures, broad-host-range promoters were diluted 1:1,000 – 1:10,000 for *Pseudoalteromonas* sp. PS5 depending on early or late stationary phase, whereas native promoters were diluted by 1:1,000. *Pseudoalteromonas* sp. PS5 with broad-host-range promoters grown in Marine Broth to stationary phase, spotted on Marine Broth agar plates as biofilms for 24h, collected and suspended in Marine Broth to an OD600 of 1.0 and diluted by 1:10,000 or 1:1,000 for strains with broad-host-range or native promoters, respectively. *Pseudoalteromonas* sp. PS5 expressing a non-luminescent plasmid (CP25-*gfp*) was used as a negative control to establish background signal from the assay. Fold changes were calculated by dividing the average RLU/$OD_{600}$ of a treatment by the background RLU/$OD_{600}$ value of another treatment.

**Transposon construction and confirmation**

The Tn10 backbone was cloned from the pLOF plasmid and includes an R6K origin of replication, an IPTG-inducible transposase, left and right inverted repeats and an ampicillin resistance gene [7]. The modular Tn10 backbone was made by PCR amplifying a section of the BTK broad host range vector (pBTK402), which contained tandem BsaI cut sites (including Type 1 and 8 overhangs), an RFP dropout reporter, and a kanamycin resistance gene. The pLOF/Km Tn10 backbone was NotI digested, cleaned up and the two parts were then Gibson assembled to generate pMMK801 (Figure S2A). To provide visual screening and fluorescent tagging capabilities, we assembled the Tn10 backbone with a CP25 promoter, a *gfp*-optim1 fluorescent reporter, and a T7 terminator (Figure S2B).

To confirm the successful integration of the Tn10 payload into *Pseudoalteromonas* sp. PS5, we selected several mutants to send for whole genome sequencing (SeqCenter). Briefly, single colonies were inoculated into 5 mL Marine Broth media and incubated overnight. 1 mL of overnight culture was spun down at 4000 g for 2 minutes, and the supernatant was removed. Genomic DNA was extracted using the Quick-DNA Fungal/Bacterial Miniprep Kit (Zymo Research) and eluted with RO water. Per SeqCenter methodology, sample libraries were prepared using the Illumina DNA Prep Kit and IDT 10 bp UDI indices, and sequenced on an Illumina NextSeq 2000, producing 2x151 bp reads. Demultiplexing, quality control, and adapter trimming was performed with bcl-convert (v3.9.3). Raw reads were uploaded to PATRIC (v3.6.12) [8, 9], and a comprehensive genome analysis was performed with default settings. Reads were trimmed with TrimGalore [10], assembled with Unicycler [11], and annotated with RastTk [12]. All strains were queried for the payload coding sequences (*gfp*, *mRuby* or *bmp2* and *KanR*).

To confirm the site of integration, we used the public server on the Galaxy Web platform to map the reads to the reference genome. Raw fastq reads were mapped to the *Pseudoalteromonas* sp. PS5 Genbank Assembly (GCF_004103255.1) using BBTools:BBMap [13] with default parameters. We reasoned that the reads containing the Tn10 inverted repeats would not map onto the genome. Therefore, we converted the unmapped FASTQ reads to FASTA format using Samtools FastX v1.13 [14]. The unmapped reads FASTA file was then made into a Blastn database in Geneious v2022.2.2, which was used to search for the Tn10 IR sequences. The unaligned portion of the reads (50-70 bp) were used to query the matching sequence on the *Pseudoalteromonas* sp. PS5 genome (Table S2).

**Coral collection and culturing**

Reproductively mature colonies of *Porites astreoides* were collected by SCUBA from the following reefs in the days before the new moon by the Mote Marine Laboratory and the Smithsonian Marine Station (SMS). Corals were collected during the 2021-2022 larval seasons [15] (June 2021 [Wonderland Reef (GPS: 24.56069, -81.50135), 21 colonies, 3222 total estimated larvae], March-April 2022 [Wonderland Reef, 10 colonies, 3735 total estimated larvae] and April-May 2022 [Oddball Reef (GPS: 24.56734, -81.45882), 15 colonies, 13,040 total larvae]) under FKNMS-2019-24 permit. Coral colonies were placed in a flow-through seawater tank at Mote's Elizabeth Moore International Center for Coral Reef Research & Restoration in Summerland Key, Florida. Larvae were collected each night for 3-4 days and were pooled for experimentation as previously described [16]. Larvae were transported to the SMS and maintained in filtered natural seawater until used in experiments.

**Coral metamorphosis assay methods**

Monospecific biofilm coral metamorphosis assays were performed as described previously [1]. Briefly, wild type and mutant strains were streaked out onto Marine Broth media and incubated overnight at 28 ºC. The next day, single colonies were inoculated into 2 mL cultures and incubated with agitation at 150 rpm for 18 hours. The optical density of the cultures was measured at 600 nm and standardized to $OD_{600}$ 0.5 (Thermo, GENESYS 180). Ceramic fragging disks (Ocean Wonders) were rinsed then sterilized by autoclave and placed into each well of a sterile, untreated 6-well plate (Falcon). Each well of the plate contained 5 mL of sterile Marine Broth and was inoculated with 100 µL of $OD_{600}$ 0.5 diluted culture. The plates were then incubated at 28 ºC for 48 hours with slow agitation (35 rpm). Control disks were either incubated for 48 hours in sterile Marine Broth media or sterile disks were added directly to the well in filtered seawater. The incubated disks were removed from the wells and rinsed under a steady stream of 0.2 µm filtered natural seawater to eliminate unattached cells. Biofilmed and control disks were randomly placed into 5 or 10 replicate deep (100 mm x 25 mm) petri dishes (Fisherbrand, FB0875711) containing 60 mL of 0.2 µm filtered seawater. Ten larvae were selected haphazardly and added to each petri dish in 10 mL, bringing the final volume of the petri dishes to 70 mL. Larvae selected for experiments were between 3 and 7 days old and were actively swimming. Figure 1D shows averaged data of 5 or 10 replicates performed on 3 separate occasions across 2 years (N=25).

**LC-MS/MS methods**

Extraction and quantification of TBP was performed as previously described [17]. Briefly, triplicate cultures of *Pseudoalteromonas* sp. PS5 were grown in 5 mL of Marine Broth media, shaking (200 rpm) overnight at 25 ºC for 16 hours and extracted with 2x volume (10 mL) of ethyl acetate. Extracts were dried under a steady stream of nitrogen gas. Dried extracts were resuspended in 100 µL methanol, filtered through a 0.2 µm centrifugal column (American Chromatography Supplies) and 10 µL was injected into a Kinetex C18 reversed-phase analytical HPLC column (5 µm, 4.6 x 150 mm, Phenomenex). TBP was quantified on an Accurate Mass QToF LC-MS/MS (Agilent 6530 Accurate Mass), run at 0.5 mL $min^{-1}$ in negative ionization mode. The same solvent system of acetonitrile and water (+0.1% formic acid (v/v)), elution profile and quantification methods were followed as previously reported [17, 18]. Synthesized TBP was used to generate a standard calibration curve as previously described [18].

**Statistics**

Data were plotted and analyzed using Prism v9 (Graphpad). Quantification of TBP was performed in 3 replicate 5 mL cultures of *Pseudoalteromonas* sp. PS5. A Shapiro-Wilk test for normality confirmed normal distribution of the wild-type strain ($P = 0.38$). Nonparametric statistics were used to confirm a significant reduction in TBP measured between the wild type and knockout strain due to unequal variances (One-tailed Mann-Whitney test, $P = 0.05$).

Coral metamorphosis assays were performed during 3 experiments over 2 years. Biofilm treatment accounted for 67.3% of total variation ($p < 0.0001$), while each experimental batch of larvae accounted for 3.1% of total variation ($p = 0.0061$) for combined coral morphogenesis (Two-way ANOVA). The response of each morphogenic phenotype (attached or unattached) was plotted (N = 25 replicate dishes, 10 larvae per dish). The data were analyzed using non-parametric statistics due to normality and variance violations for the Δ*bmp2* treatment (Shapiro-Wilk test). Kruskal-Wallis tests determined that the medians varied significantly ($P < 0.0001$), while Dunn’s multiple comparisons tests were used to compare the anticipated differences in the sum of ranks between the wildtype, Δ*bmp2* biofilm and control treatments for attached and unattached metamorphosis independently, as well as combined morphogenesis. P-values are reported as multiplicity adjusted p-values to account for multiplicity of comparison.

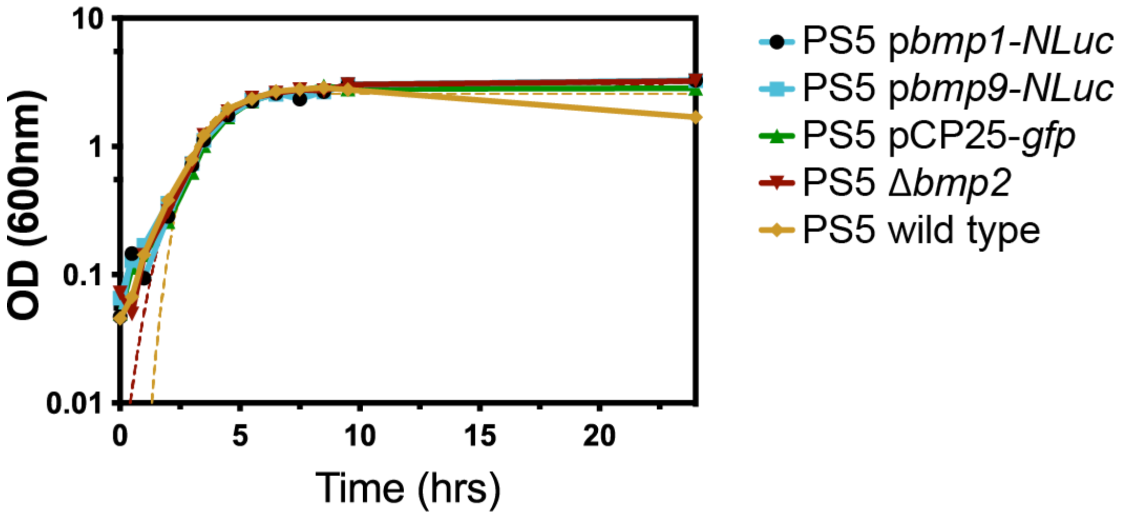


**Figure S1: Growth curve of *Pseudoalteromonas* sp. PS5 strains.** Growth curve of *Pseudoalteromonas* sp. PS5 wild type, Δ*bmp2* and strains expressing plasmids *bmp1-NLuc*, *bmp9-NLuc,* and CP25-*gfp*. Optical density (OD) measurements were taken at 600 nm wavelength and graphed on a $Log_{10}$ scale. Dotted lines correspond to wild type and Δ*bmp2* Gompertz non-linear regression fit. Plotted is the average of two biological replicates.

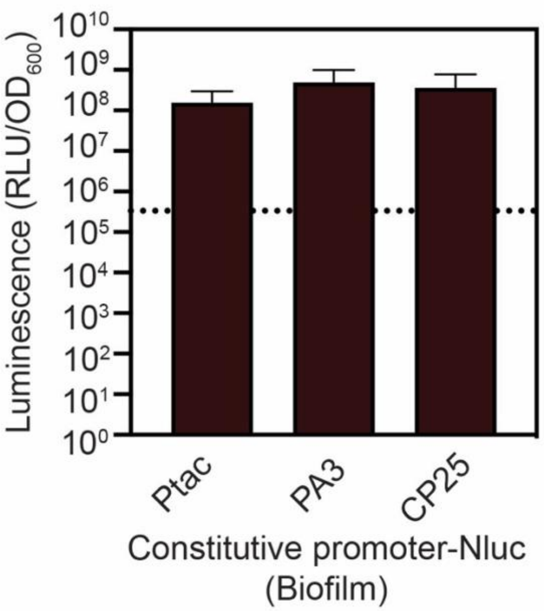


**Figure S2. Quantification of broad-host-range promoter expression in *Pseudoalteromonas* sp. PS5.** Modular broad-host-range promoter parts, including the hybrid Ptac promoter [6] and the synthetic CP25 promoter, were cloned to drive NanoLuciferase expression in *Pseudoalteromonas* sp. PS5 [5]. Error bars show standard deviation of the average of N= 4 biological replicates. The dotted line (Y= 333,700 RLU/$OD_{600}$) represents the average signal of the non-luminescent negative control treatment (*Pseudoalteromonas* sp. PS5 expressing CP25-*gfp*).

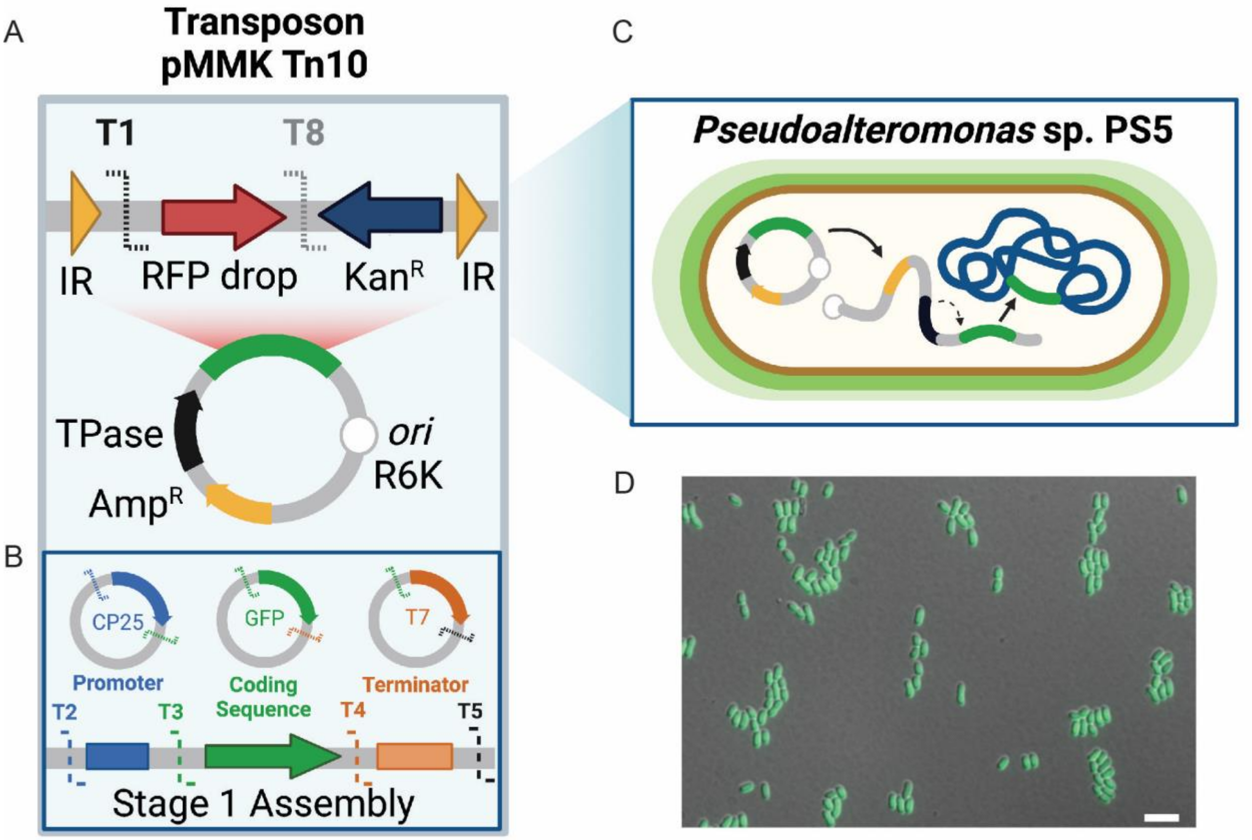


**Figure S3. Design and construction of a transposon backbone for integrated tagging and complementation in *Pseudoalteromonas* sp. PS5.** (A) A Type 8 transposon backbone was constructed in this study to enable integrative tagging and complementation options compatible with the Marine Modification [6] and Bee Toolkit [5]. The backbone contains an R6K origin of replication, a *bla* resistance gene, and an IPTG-inducible transposase. The resulting Type 8 part, pMMK801, adds an RFP dropout, flanking BsaI sites and a kanamycin resistance gene cloned in between the inverted repeats (IR). (B) To confirm integration, we performed a Stage 1 Golden Gate Assembly with a previously successful expression payload (CP25-*gfp*-T7) [6]. (C) We successfully conjugated the transposon into *Pseudoalteromonas* sp. PS5 and (D) observed uniform fluorescence in the transconjugants using fluorescence microscopy. Scale bar is 10 µm.

**Table S1. Genomic insertion loci for Tn10 mutants in *Pseudoalteromonas* sp. PS5**

| Mut | Genotype | Mapped PS5 WGS contig | Insertion locus | Gene Function | Accession |
|---|---|---|---|---|---|
| 1 | PS5:Tn10-Ptac-*gfp*-T7 | NZ_RCSQ01000015.1 | 8938 | DUF4856 domain-containing protein | WP_128729848.1 |
| 8 | PS5:Tn10-CP25-*gfp*-T7 | NZ_RCSQ01000092.1 | 8552 | Hypothetical protein | WP_128731616.1 |
| 6 | PS5:Tn10-CP25-*mRuby*-T7 | NZ_RCSQ01000098.1 | 9251 | Non-ribosomal peptide synthetase | WP_128731684.1 |

**Table S2. List of strains used in this study**

| Strain no. | Strain | Genotype | Source |
|---|---|---|---|
| NJS595 | *Pseudoalteromonas* sp. PS5 | Wild type | [1] |
| NJS597 | *Pseudoalteromonas* sp. PS5 | StrR | [6] |
| NJS602 | *Pseudoalteromonas* sp. PS5 Δ*bmp2* | StrR Δ*bmp2* | This Study |
| NJS712 | *Pseudoalteromonas* sp. PS5 Tn10::Ptac-*gfp*-T7 | StrR KanR | This Study |
| NJS726 | *Pseudoalteromonas* sp. PS5 Tn10::CP25-*gfp*-T7 | StrR KanR | This Study |
| pNJS488 | *Escherichia coli* S17-1 | TpR SmR recA thi pro (rK− mK+) RP4: 2-Tc:Mu: Km Tn7 λpir | [19] |
| NJS604 | *Escherichia coli* MFDpir | MG1655 RP4-2-Tc::[ΔMu1::aac(3)IV-ΔaphA-Δnic35-ΔMu2::zeo] ΔdapA::(erm-pir) ΔrecA | [20] |
| pNJS033 | *Escherichia coli* SM10 | thi thr leu tonA lacY supE recA::RP4-2-Tc::Mu Km λpir | [21] |

**Table S3. List of plasmids used in this study**

| Plasmid | Type | 5' Site | 3' Site | Description | Marker | Origin | Source |
|---|---|---|---|---|---|---|---|
| pBTK001 | Entry vector | N/A | N/A | Entry vector for generating new parts | CamR | p15A | [5] |
| pYTK008 | Connector | 1 | 1 | ConLS' connector | CamR | ColE1 | [22] |
| pBTK107 | Promoter | 2 | 2 | CP25 synthetic promoter, RBS | CamR | ColE1 | [5] |
| pBTK121 | Promoter | 2 | 2 | PA3 strong early promoter (phage), RBS | CamR | ColE1 | [5] |
| pMMK201 | Promoter | 2 | 2 | ptac hybrid promoter, RBS | CamR | ColE1 | [6] |
| pMMK204 | Promoter | 2 | 2 | PS5 bmp1, RBS | CamR | ColE1 | This study |
| pMMK205 | Promoter | 2 | 2 | PS5 bmp9, RBS | CamR | ColE1 | This study |
| pYTK047 | GFP Dropout | 2 | 4 | GFP dropout (internal BsaI sites) | CamR | ColE1 | [22] |
| pBTK205 | Coding sequence | 3 | 3 | GFP optim-1 | CamR | ColE1 | [5] |
| pBTK206 | Coding sequence | 3 | 3 | NanoLuciferase | CamR | ColE1 | [5] |
| pYTK034 | Coding sequence | 3 | 3 | mRuby2 | CamR | ColE2 | [22] |
| pBTK305 | Terminator | 4 | 4 | T7 terminator | CamR | ColE1 | [5] |
| pYTK073 | Connector | 5 | 5 | ConRE' connector | CamR | ColE1 | [22] |
| pBTK402 | Origin, Marker | 8 | 8 | mRFP1 dropout, pMMB67EH derivative | KanR | RSF1010 | [5] |
| pLOFkm | Transposon | 8 | 8 | Tn10 transposon | KanR, AmpR | R6k | [7] |
| pMMK801 | Origin, Marker | 8 | 8 | Tn10 transposon, mRFP1 dropout | KanR, AmpR | R6k | This study |
| pMMK802 | Stage 1 Dropout | 2 | 4 | Tn10 GFP dropout | KanR, AmpR | R6k | This study |
| pMMK806 | Stage 1 assembly | 1 | 5 | p402-bmp1p-NLuc-T7 | KanR | RSF1010 | This study |
| pMMK807 | Stage 1 assembly | 1 | 5 | p402-bmp9p-NLuc-T7 | KanR | RSF1010 | This study |
| pMMK803 | Stage 1 assembly | 1 | 5 | Tn10-CP25-GFP-T7 | KanR, AmpR | R6k | This study |
| pMMK804 | Stage 1 assembly | 1 | 5 | Tn10-CP25-mRuby-T7 | KanR, AmpR | R6k | This study |
| pMMK805 | Stage 1 assembly | 1 | 5 | Tn10-Ptac-gfp-T7 | KanR, AmpR | R6k | This study |
| pMMK809 | Stage 1 assembly | 1 | 5 | p402-PA3-NLuc-T7 | KanR | RSF1010 | [5] |

| | | | | | | | |
|---|---|---|---|---|---|---|---|
| pMMK810 | Stage 1 assembly | 1 | 5 | p402-CP25-NLuc-T7 | KanR | RSF1010 | [5] |
| pMMK811 | Stage 1 assembly | 1 | 5 | p402-Ptac-NLuc-T7 | KanR | RSF1010 | [6] |
| pCVD443 | Suicide vector | N/A | N/A | sacB, pGP704 derivative | AmpR | R6k | [4] |
| p443bmp2 | Suicide vector | N/A | N/A | pCVD443::Δ*bmp2* | KanR, AmpR | R6k | This study |

**Table S4. List of Primers used in this study**

| Primer | Sequence |
|---|---|
| p443_PS5_dbmp2_A1 | GGTTAAAAAGGATCGATCCTCTAGACGAACCACCACATTCTCCTT |
| p443_PS5_dbmp2_B1 | TGGGTAATTCCCTTAACTTGCGTTCATTACATTGCCACCTTATTTA |
| p443_PS5_dbmp2_C1 | TAAATAAGGTGGCAATGTAATGAACGCAAGTTAAGGGAATTACCCA |
| p443_PS5_dbmp2_D1 | CAACGTGAATTCAAAGGGAGAGCTCACGGCATGACTTGTCTACCC |
| PS5_dbmp2_seq_F1 | CACCATTGCTTGAACTTGGT |
| PS5_dbmp2_seq_F2 | AGGCTTTGGTTTGGTTGATG |
| PS5_dbmp2_seq_F3 | AGCAGAAGCAGGTTCCGATA |
| PS5_dbmp2_seq_F4 | ATCGCCGATATGGAAGTGAG |
| PS5_dbmp2_seq_R1 | TGTGCCTCATTCCATTCAAA |
| PS5_dbmp2_seq_R2 | CTGCCATTGGTCACATAGGA |
| PS5_dbmp2_seq_R3 | TGCCTTTGACTCGTAGCGTA |
| pYTK034_vector_amp_F | ATCCTGAGACCAGACCAATAAAA |
| pYTK034_vector_amp_R | ATGAGACCGACTACGGTTATCC |
| pYTK034_PS5bmp2_gbsn_F | GGATAACCGTAGTCGGTCTCATATGAACGGATTTACACATTATGACG |
| pYTK034_PS5bmp2_gbsn_R | TTTTATTGGTCTGGTCTCAGGATTTAACTTGCCATTTGTTTACGG |
| PS5_bmp2_R1 | GCATCCATATCCTCCGCTAA |
| p107_bbamp_F | TATGTGAGACCAGACCAATAAAAA |
| p107_bbamp_R | CGTTTGAGACCGACTACGGTTA |
| pBTK107_PS5bmp1_promoter_gbsn_F1 | ACCGTAGTCGGTCTCAAACGCGAACCACCACATTCTCCTT |
| pBTK107_PS5bmp1_promoter_gbsn_R1 | GTTTTTTATTGGTCTGGTCTCACATAGCAGCACCTTCGAGTAGATCG |
| pBTK107_PS5_bmp9_fusion_gbsn_F1 | ACCGTAGTCGGTCTCAAACGCAGGGATTTTTGCACCGTAA |
| pBTK107_PS5_bmp9_fusion_gbsn_R1 | GTTTTTTATTGGTCTGGTCTCACATAACTTCGGTCTCGATGGTTT |
| ps5bmp1_seq_f | CCACCACATTCTCCTTCAATAC |
| ps5_bmp89_seq_f | GCACCTTCGAGTAGATCGTTATT |
| pLOF_NotI_Kan_F | GGTCAGCCTGAATACGCGTGCGGCCGCGAGCACCTGGTCGCTTTC |
| pLOF_NotI_Kan_R | ACGCGTGGCCGCCTAGGCCGCGGCCGCCGGGCAAGTACGACATCAC |
| pTn10_Transposase_F1 | CACACATGGTTACGCTTTGG |
| pTnt10_TcR_R1 | GGCCCATTACTGTTTGCTGT |
